# Dissecting PCNA function with a systematically designed mutation library in yeast

**DOI:** 10.1101/352286

**Authors:** Qingwen Jiang, Weimin Zhang, Chenghao Liu, Yicong Lin, Qingyu Wu, Junbiao Dai

## Abstract

Proliferating cell nuclear antigen (PCNA), encoded by *POL30* in *Saccharomyces cerevisiae*, is a key component of DNA metabolism. Here a library consisting of 308 PCNA mutants was designed and synthesized to probe the contribution of each residue to its biological function. Five regions were identified with elevated sensitivity to DNA damaging reagents using high-throughput phenotype screening. Using a series of genetic and biochemical analyses, we demonstrated that one particular mutant, K168A, which displayed severe DNA damage sensitivity, abolished the DNA damage tolerance (DDT) pathway by disrupting interactions between PCNA and Rad5p. Subsequent domain analysis showed that the PCNA/Rad5p interaction is prerequisite for the function of Rad5p in DDT. Our study not only provides a resource in the form of a library of versatile mutants to study PCNA functions, but also reveals a key regulatory function of Rad5p, which highlights the importance of the PCNA-Rad5p interaction.

**Author summary:** PCNA is regarded as the maestro of DNA replication fork because of the astonishing ability to interact with lots of partner proteins that participate in various DNA metabolism processes. However, it has remained elusive as to how does PCNA orchestrate these functions in harmony. Here, we constructed a systematic mutation library of PCNA, which covers every amino acid to map the functional sites of it. This carefully designed synthetic mutant pool could be generally useful and serve as a flexible resource, such as dissecting the functional mechanism of PCNA by genetic relationship analysis with key proteins through Synthetic genetic array. We further dissected the intrinsic mechanism for damage sensitivity of PCNA^K168A^, the most severe DNA damage sensitive mutant in our alanine scanning mutation library, this helps us to get better understanding of how PCNA participates in DNA damage tolerance (DDT) pathways. Our findings indicate that K168 site is vital for the interaction between DDT related partner proteins and PCNA, and also highlight the importance of the PCNA-Rad5p interaction.

## Introduction

Accurate and efficient propagation of genetic information to descendants is the central task for dividing cells. Nevertheless, DNA is highly vulnerable to many types of genotoxic challenges that can lead to DNA damage and replication fork stall, termed DNA replication stress (1). Inappropriate molecular management of replication stress may result in genetic variation, replication fork collapse or cell death. Proliferating cell nuclear antigen (PCNA) plays central roles in DNA replication and repair by guaranteeing replisome fluidity (2). This toroid shaped homotrimer is structurally superimposable across eukaryotes consistent with evolutionary conservation of their role as sliding clamps (3). Based on a common mode of action, PCNA was reported to interact with various partners to orchestrate different DNA metabolic functions in harmony (3, 4).

Several established mechanisms of DNA damage tolerance (DDT), which are vital for relieving replication fork stalling and ensuring complete duplication of the genome even in the presence of bulky DNA lesions, are regulated by distinct post-translational modifications (PTMs) of PCNA (3, 5, 6). Monoubiquitylation of PCNA on K164 by the Rad6-Rad18 complex triggers an error-prone bypass mechanism through the recruitment of dedicated translesion polymerases. These enzymes can replicate across lesions (translesion synthesis; TLS) (7). Polyubiquitylation of PCNA on K164 site by Mms2/Ubc13-Rad5 complex is involved in a recombination-related mechanism, which can switch replication from the damaged site to the undamaged sister chromatin as template, which is thus named the template switching pathway (TS) (8). A typical symbol of this pathway is sister chromosomal junction (SCJ) shaped like an “X molecule” similar to the Holliday junction (9). Several important genes involved in this pathway have been identified by detecting the presence of SCJs (10, 11), leading to a “sketch map” of this pathway. Another type of modification on PCNA is SUMOylation at lysines 127 and 164 by Ubc9 or Siz1, which acts as inhibitors of homologous recombination by recruiting Srs2p to remove Rad51p from chromatin, and thus keeps certain potentially deleterious HR pathways in check (12).

Although PCNA modifications by SUMO and ubiquitin are known to be crucial for DDT, the regulation or choice among these three pathways remains poorly understood. Structural analysis of the PCNA -Siz1-Srs2 and PCNA-polη complexes have shed some light on the working mechanism of HR and TLS pathways (13, 14), whereas the TS pathway remain largely elusive. It is unclear how Rad5p, a key component which functions both upstream, as a ubiquitin ligase, and downstream, as a helicase lacking a canonical PIP (PCNA-interacting peptide)-box, participates in this pathway (11). In addition, it is also unclear how polyubiquitination on PCNA is correlated with the TS pathway, and why some well-known HR-related proteins are involved in TS as reported in recent years (10).

On the other hand, not all functions of PCNA rely on their modification states and indeed, previous mutagenesis studies have identified many unmodifiable residues which are crucial for PCNA to participate in DNA replication, repair (15–20), chromatin dynamics and cell cycle regulation (21, 22). Functional regions such as the inter-domain connecting loop (IDCL) (16), trimer interface (23–25) and the DNA interface helix were identified, which lead to the present model of the exchange of PCNA partners and the loading and sliding mechanism of the PCNA clamp. Currently, mutational analyses of PCNA structure-function relationships are limited to defined subsets of residues (26).

In this study, we generated a PCNA mutant library consisting of 308 alleles to cover every residue within this protein based on a carefully designed synthetic construct, which could be generally useful and serve as a flexible resource. We systematically substituted each residue with alanine, while replacing each native alanine residue with serine. At the IDCL region, a more comprehensive mutagenesis strategy was applied, including swapping the charge status of each residue and substituting each modifiable residue with another amino acid residue to mimick either modified or unmodified status, when possible. We screened the library for individual mutants that made yeast fail to resist DNA damaging agents such as methyl-methanesulfonate (MMS) and hydroxyurea (HU), which revealed several important regions within PCNA. We then focused our study on one particular mutant, K168A, which showed severe sensitivity to DNA lesions and revealed the importance and the intrinsic acting mechanism of this site for PCNA during DDT.

## Results

### The design of a synthetic PCNA cassette

To generate a PCNA mutant library that could serve as a useful resource for various research purposes, we carefully designed a synthetic base construct (Fig 1A) according to the following rules: 1) Only synonymous changes were incorporated into the synthetic coding sequence (named synPCNA) to preserve the identical amino acid sequence as that of the wild type PCNA (wtPCNA), i.e. *POL30* in *S. cerevisiae*. No more than five consecutive nucleotides remain the same between synPCNA and wtPCNA (Fig S1A), largely eliminating the possibility of recombination between them; 2) The native 3’ untranslated region (3’ UTR) was replaced by the one from *ADH1*; 3) A molecule barcode (named TAG) at the length of 20-nucleotides, adopted from the yeast knockout library (27), was inserted downstream of the 3’ UTR, which could serve as a unique ID for each construct; 4) The *LEU2* gene (ChrIII 90972-92719) flanked by two loxP sites was placed adjacent to the TAG, allowing selection of the synthetic construct and marker-swapping, if necessary; 5) The abovementioned parts are flanked by the native 5’ and 3’ UTR of PCNA at the length of more than 500 base pair (bp) each, which not only allow the entire construct to be integrated at the native locus, but also put the synPCNA under the control of the native *POL30* promoter; 6) The designed cassette was chemically synthesized and cloned into pRS414, which could be either used as an episomal plasmid or released by restriction enzyme *Ear*I for integrating into the endogenous *POL30* locus.

**Fig 1.**
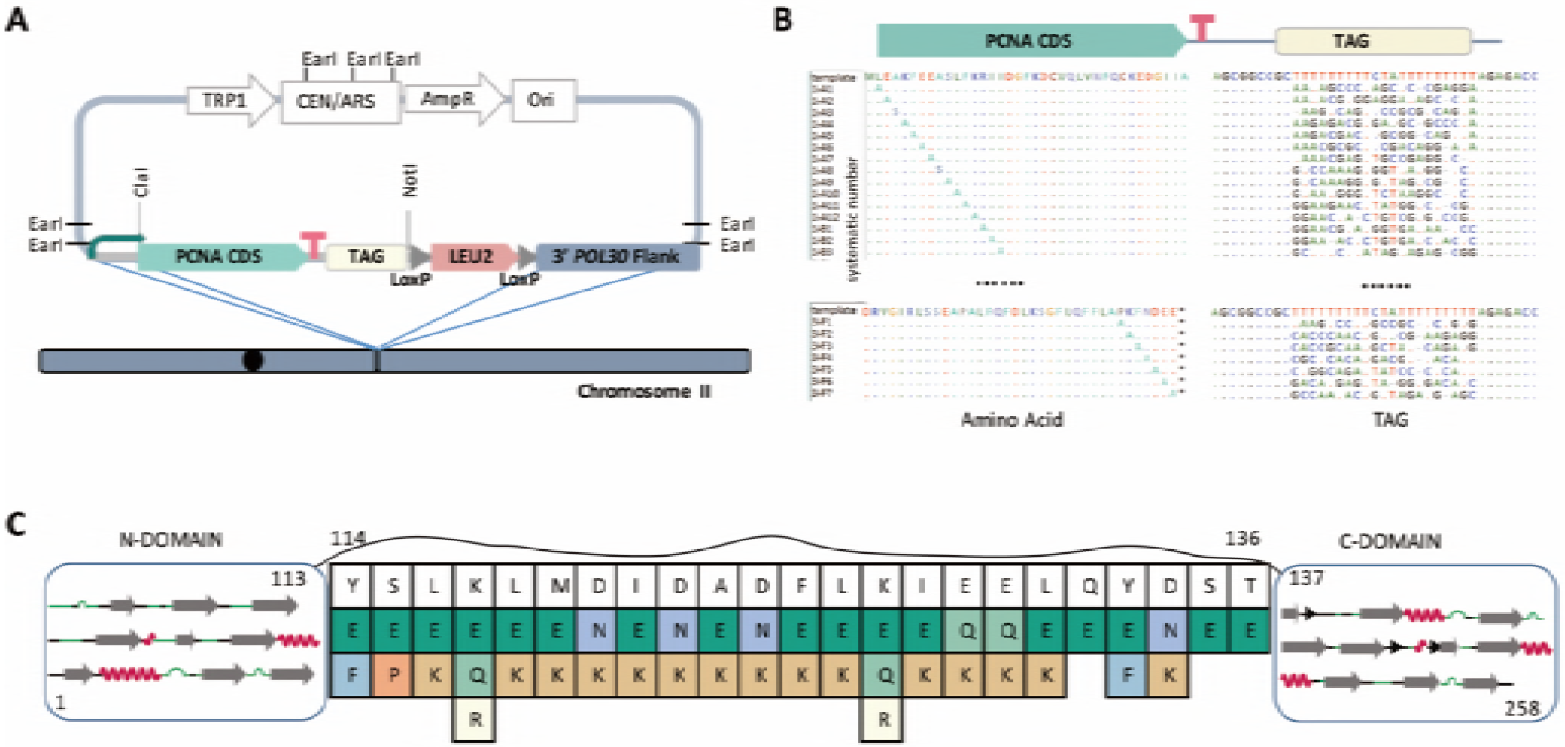
The Design of PCNA mutant Library. **(A)** Schematic representation of the synthetic PCNA cassette within the *pRS414* vector. **(B)** The design of systematic Alanine mutagenesis strategy. The tag sequence for each mutant is different, which are listed in table S2. **(C)** Illustration of versatile mutations at the IDCL region. The secondary structure of N- and C-terminal domains of PCNA is shown. The original amino acid residues are shown on the top. The colored residues represent the amino acids to replace the native ones respectively.

We at first tested if synPCNA could replace the native PCNA by plasmid shuffling assay. A strain was constructed by integrating synPCNA at the endogenous *POL30* locus and presenting a copy of the wtPCNA in a centromeric plasmid with URA3 marker. As shown in Fig S1B, cells with only synPCNA grew similarly to the wild type, indicating that synPCNA could carry out the essential function of its wild type equivalent. To further probe if there were any subtle differences between the two versions of PCNA, a serial dilutions dot assay was carried out to monitor cell fitness under various growth conditions, which showed that the strain containing synPCNA was indistinguishable from the wild type (Fig S1B). Additionally, the expression of synPCNA was also examined by western blot, which showed no difference between the synthetic and native PCNA (Fig S1C). We conclude that synPCNA is fully functional, at least under the conditions tested.

### A versatile library of synthetic PCNA mutants

Based on synPCNA, two major classes of systematic mutants were designed. The first class consists of 257 variants, including an alanine substitution of each residue. Each native alanine residue was replaced by serine (Fig 1B). The second class focused on residues at the IDCL region, which is key for interacting with “PIP” box containing partner proteins (Fig 1C). These mutants were generated following the rationale listed in table S1. For example, lysine residues were mutated to arginine and glutamine to potentially mimic constitutively deacetylated/acetylated states and to glutamic acid to swap the charge state. In addition, three mutations of Y211 were constructed to mimic constitutive phosphorylation or dephosphorylation states (28, 29) since this site is known to be phosphorylated in human. At last, two previously reported PCNA mutants, G178S and LI126, 128AA (17, 24), were also constructed to serve as control for different assays.

In addition of changes at the designated amino acid residue, a unique 20 bp TAG sequence was assigned to each mutant (Table S2), which could enable us to pool all the mutants and perform complex assays such as competitive growth in a chemostat (30).

All 308 mutants were synthesized and supplied as individual clones in bacteria. The arrangement of each mutant is shown in Fig S2. The plasmids were prepared, linearized and transformed into strain QJY001, in which the chromosomal *POL30* gene had been replaced by KanMX4 and a copy of wild-type *POL30* was supplied on a *URA3*-marked centromeric plasmid. After the mutant construct was integrated, the wild type plasmid was removed by plating the cells in medium containing 5-fluoroorotic acid (5-FOA), which eventually produced a library of yeast strains each containing only a copy of the mutant construct integrated at its endogenous locus. Two independent genotypic correct yeast colonies for each mutant were isolated and arrayed in 96-well plate for stock as listed in Fig S3.

### High-Throughput phenotyping of individual PCNA mutants

All viable yeast strains containing individual PCNA mutant were subjected to various growth tests including sensitivity to different temperatures (16°C, 30°C, 37°C and 39°C), DNA damage stresses (Hydroxyurea (HU), Methyl methanesulfonate (MMS), Camptothecin (CPT) and various dosage of Ultraviolet (UV) irradiation) and microtubule disruption stress (benomyl).

We found most of the mutants showed no obvious defects under above tests and, as expected, most of the phenotypes result from treatment with DNA damaging reagents (Table S3). Among the alanine scanning mutants, 51 out of 257 (19.8%) mutants showed sensitivity to at least one DNA damaging reagent. This finding is consistent with the biological function of PCNA, which is tightly involved in DNA replication and repair (4). Therefore, we focused on dissecting the underlining mechanisms of DNA damage sensitivities among these mutants in the remaining studies.

From the alanine-substitution mutants, we identified three major classes of mutants which were sensitive to MMS (94%), HU (20%) and UV irradiation (20%) respectively. As shown in the Venn diagram (Fig 2A), 36 mutants were only sensitive to MMS, 2 mutants showed only sensitivity to HU while 10 mutants showed sensitivity to UV also had sensitivity to either HU or MMS or both. There were 4 mutants showing sensitivity to all three kinds of DNA damage stress. We calculated the evolutionary conservation value of these amino acids that were sensitive to DNA damage, if mutated, and found they were more conserved comparing with total amino acid residues (Fig 2B), revealing that they are more important. The DNA damage sensitivity scores of all the mutant alleles are listed in table S3.

**Fig 2.**
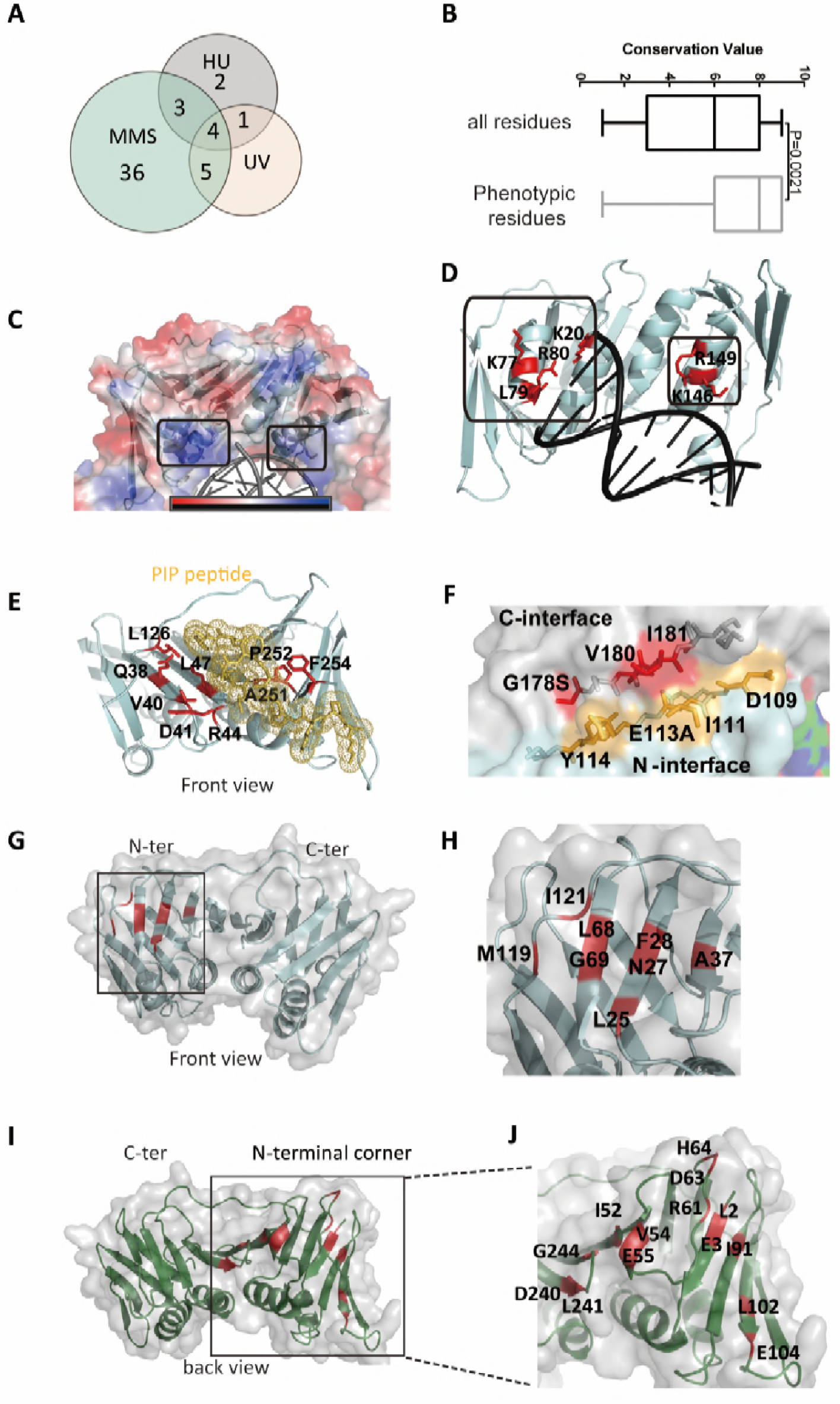
Identification of PCNA mutants with increased sensitivity to DNA damage reagents. **(A**) Venn diagrams showing the overlaps of mutants. **(B)** Correlation between damage-sensitive alleles and evolutionary conservation. Evolutionary conservation scores were calculated using ConSurf. Whiskers in the box and whiskers plot represent minimum and maximum values excluding outliers, defined as 3/2 times higher (or lower) quartile. **(C-J)** Positions of mutated residues within the PCNA crystal structure. The identified residuals were marked in red (red and yellow in (f)) and labeled. **(C)** Amino acids at inner helix region are marked by square on PCNA electric potential map. **(D)** Enlarged map for indicating the residues of (C). **(E)** PIPbox peptide interaction sites mapped on complex structure of PCNA and CDC9 generated PIP peptide. PCNA is in cyan and PIP peptide is in yellow. **(F)** Phenotypic sites located at trimer interface. Red: the C terminal sites; Yellow: the N terminal sites. **(G)** Phenotypic sites located at N terminal front region. **(H)** Detail annotation of (G). **(I)** Phenotypic sites located at N terminal back region. **(J)** Detail annotation of (I).

When mapping these sites to the crystal structure of the PCNA trimer, we identified five structural coherent clusters. The first cluster is located at the central alpha helix which is close to the DNA strand and forms a positive charge cluster (Fig 2C). According to previous studies, these positive charges may facilitate PCNA binding to negatively charged DNA (31) (Fig 2D). The second cluster is the PIPbox peptide binding region (Fig 2E). By mapping the sites in the crystal structure of the Cdc9 PIPbox peptide in complex with PCNA (PDB: 2OD8) (32), we showed that the sensitive sites (highlighted in red) are important for interacting with the PIPbox peptide. The third cluster lies on the interface between two monomers (Fig 2F). Mutations at these sites might weaken the formation of the PCNA trimer. Furthermore, we identified two regions at the N terminal domain: The N terminal front region (Fig 2G-H) and the N terminal back region (Fig 2I-J). The phenotypic sites on the front region just buried under the IDCL loop and this cluster may act as interaction hub for some partner proteins. Other novel sites which were not observed in previous studies were located at the back side of the N terminal domain. This cluster may also act as a face for interaction with important partner molecules. Interestingly, relatively fewer mutations (K164A, K168A, K196A) in the C terminal domain corner showed sensitivity to DNA damage stress. Since the K196 site and K164 site have been well studied previously and the K168 site is adjacent to K164 site, we did not show these sites as a functional cluster in this figure.

As a conclusion, we systematically mapped the functional sites of PCNA which not only fleshed out the functional region reported in previous studies, but also identified a new cluster located on the N terminal domain.

### The side chain of PCNA K168 is crucial for DDT and the deficiency of K168A is distinct from K164 site modification

Next, we focused on one mutant, K168A, which is severely sensitive to UV, HU and MMS (Table S3). We confirmed its damage sensitivity by growth assay in rich medium with or without MMS. The K168A mutant grew comparatively to that of wild type in medium without MMS (doubling time (DT) at 1.61hr and 1.68hr respectively). In contrast, it grew much more slowly in the presence of 0.005% MMS (DT=2.2hr, Fig 3A). According to the results from FACS analysis (Fig 3B), the longer doubling time comes from a retarded cell cycle in K168A after MMS treatment. Nevertheless, K168A showed identical cell cycle progression to that of wild type in the absence of MMS (Fig S4).

**Fig 3.**
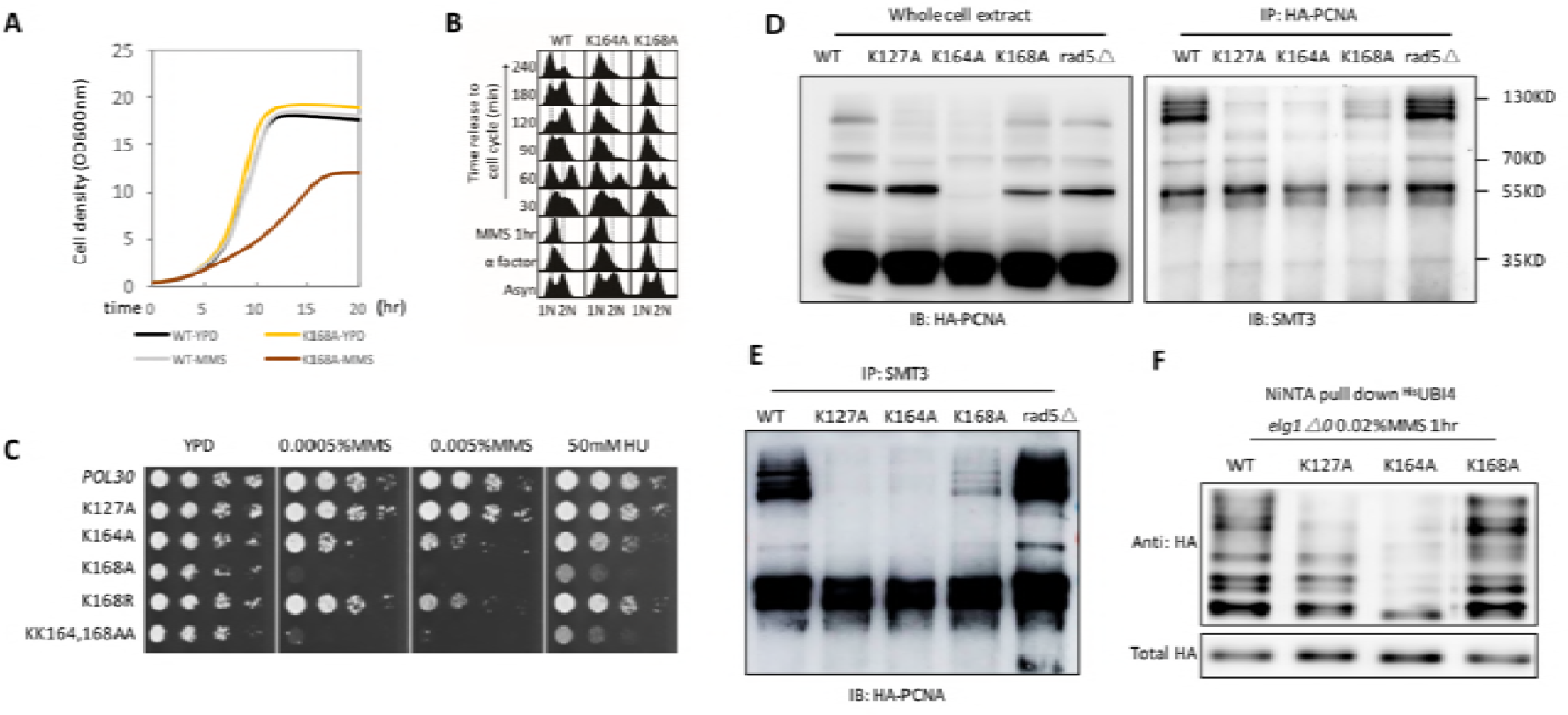
The side chain of PCNA K168 is crucial for DDT and the deficiency of K168A is distinct to K164 site modification. **(A**) Growth curves of the wild-type and K168A mutant in YPD or YPD containing 0.005%MMS. **(B)** K168A mutant cells are blocked in G1-S phase after transient expose to MMS. Cells were synchronized by α-factor for 2 hrs, exposed to MMS for 1 hr and released to cell cycle in YPD after three times wash. Samples were taken at different time points and processed for cell cycle analyzed by FACS. **(C)** Damage-sensitivity test of PCNA alleles. **(D)** Analysis of PCNA SUMOylation. HA-tagged PCNA alleles were blotted with anti-HA antibody. The modification band are labeled aside. **(E)** Analysis of PCNA ubiquitination. The same strains in (D) were used. The ubiquitinated proteins were enriched by pulling down the His-tagged Ubi4p, and then blotted with anti-HA antibody.

Since lysine is one of the major target of post-translational modifications (PTMs), we at first tested whether K168 can be modified. As a result, we failed to detect any modifications at this site using the purified PCNA from yeast by mass spectrometry. Next, we asked whether the arginine substitution, which blocks both ubiquitination and SUMOylation but keeps a similar side chain charge, displays similar phenotypes as K168A. Interestingly, the K168R mutant strain didn’t show significant sensitivity to DNA damage (Fig 3C), ruling out the possibility that the defect comes from the loss of ubiquitination or SUMOylation on K168 and hinting that the presence of positively charged side chain at this position might be critical. Since K168 is near to K164, a site that can be modified with either SUMOylation or ubiquitination and participates in three different repair pathways (12, 33, 34), we hypothesized that the side chain of K168 may warrant site specificity of K164 during PTM and K168A mutation might compromise normal modifications at K164. To detect SUMOylated PCNA, we treated cells with MMS for 2hr to accumulation the modification and immunopreciped the HA tagged PCNA with HA antibody, then sumoylated PCNA were immunoblotted with SMT3 antibody (Fig 3D) under denatured condition. The Western Blot shows that the alanine substitution at K168 resulted in a decreased amount of PCNA SUMOylation (Fig 3D). Similarly, immunoprecipitate with SMT3 antibody and immunoblot with HA antibody reveal the same conclusion (Fig 3E). Consistently, K168R did not affect PCNA SUMOylation (Fig S5). It is known that SUMOylation could prevent recombination (12), so decreased SUMOylation level may lead to increased HR. The released HR is actually a salvage repair pathway for DNA damage, because both *siz1△* and *srs2△* can rescue the damage sensitivity caused by TLS and TS pathway deficiency (12). Therefore, the decreased K164-SUMO is not directly responsible for, but may contribute to, the extreme damage sensitivity of K168A mutant.

Next, we tested whether PCNA ubiquitination is also affected by K168A. To enrich the ubiquitinated PCNA, we used His-tagged Ubi4 to pull down the modified proteins and specifically detected PCNA in the enriched samples. As shown in Fig 3F, unlike K127A or K164A mutant in which ubiquitylated PCNA isoforms were largely reduced, both the modified protein amount and the band distribution pattern in K168A were very similar to that of wild type (Fig 3F), suggesting that K168A mutation is not likely to interfere ubiquitination of PCNA. In addition, we examined the location of K168 in the crystal structure of PCNA with mono-ubiquitin (7), we found the ubiquitin is far away from K168. Together, these results suggest that the loss of side chain at K168 does not affect PCNA ubiquitylation but will lead to the reduction of PCNA SUMOylation.

### K168A mutant blocks interaction between PCNA and Rad5p

Since HR can rescue the damage sensitivity caused by deficiency in TLS or TS pathway, we tested whether deleting *SIZ1* or *SRS2*, which could completely release the SUMOylation-inhibited HR, can also rescue the K168A mutant. As shown in Fig 4A, both *siz1△* and *srs2Δ* can rescue damage sensitivity of K168A. This result further supported that decreased SUMOylation is not the reason for DNA damage sensitivity of K168A.

**Fig 4.**
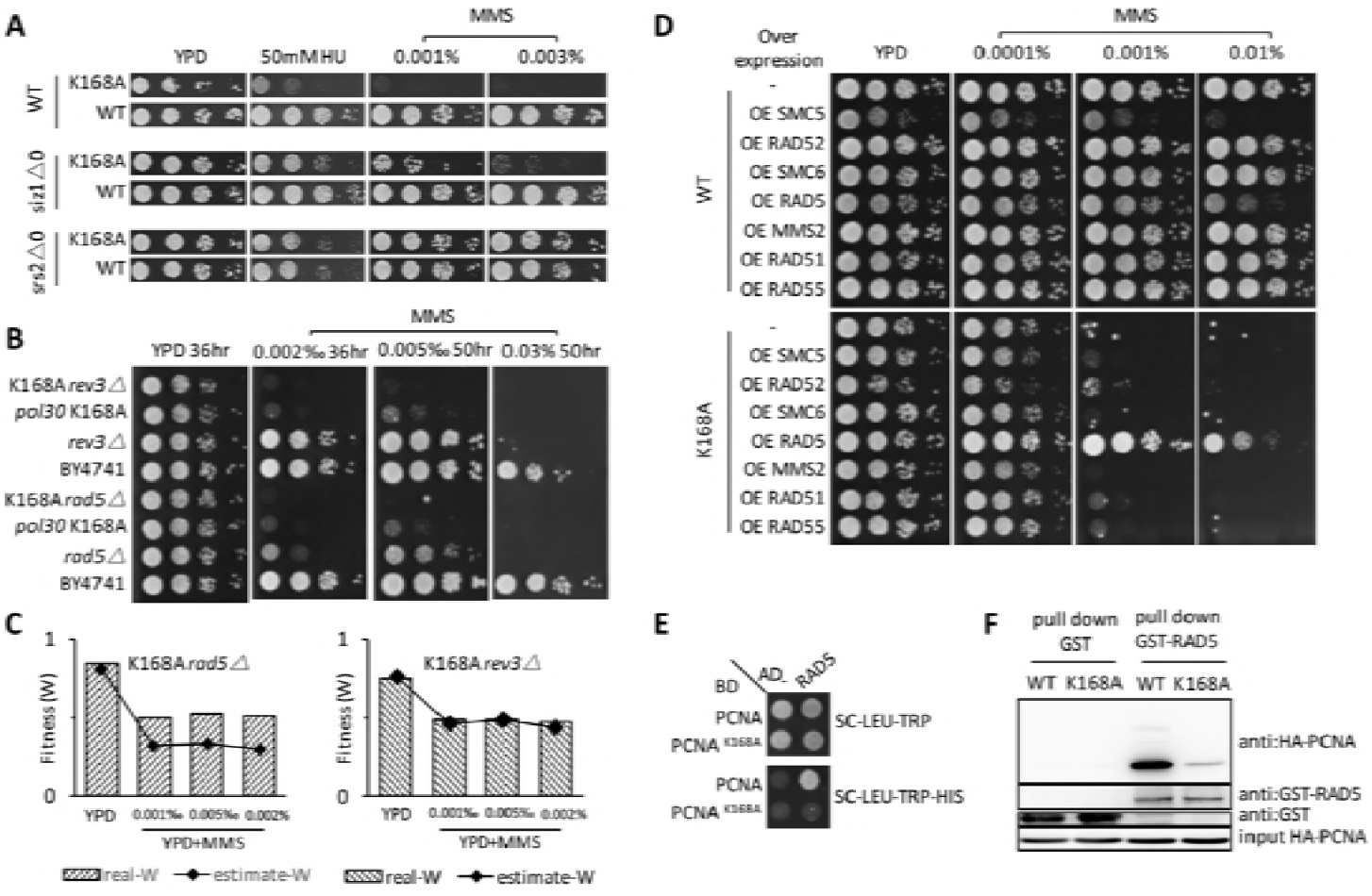
K168A reduces interactions between PCNA and Rad5p. **(A)** Genetic interactions between Srs2 and the PCNA alleles. **(B)** Genetic interactions between *REV3 RAD5* and K168A. **(C)** Classification of genetic interaction with estimated fitness (W) and measured fitness of double mutant. The estimated fitness was calculated based on fitness of single mutation fitness. **(D)** Overexpression of TS related gene in PCNA alleles. The indicated gene was over expressed by knock in a *GPD* promoter by standard tool box. **(E)** Interaction between Rad5p and PCNA K168A mutant measured by yeast two-hybrid assay **(F)** Interaction between Rad5p and PCNA K168A mutant measured by co-immunoprecipitation analysis. GST-tagged Rad5p was expressed in strain harboring HA-tagged PCNA alleles.

Considering that *siz1△* or *srs2△* can rescue TLS or TS pathway deficiency, we further tested whether the damage sensitivity of K168A mutant is caused by deficiency in these two pathways. We then checked whether the TLS pathway was affected by mutating *REV3*, a key component in TLS pathway, in wild-type or K168A strain and tested the damage sensitivity of single and double mutants (Fig 4B). We found the disruption of TLS pathway enhanced the sensitivity of K168A to the damaging reagent, suggesting the hypersensitivity of K168A is probably not caused by TLS pathway defection.

Finally, we examined whether the template switching (TS) pathway was involved. We found that knocking out polyubiquitylation-related E3 ligase *RAD5* resulted in severe sensitivity (Fig 4B). By greatly reducing the MMS concentration in the assay (0.0002%), we were able to detect the damage sensitivity of *rad5△* and K168A mutants and the double mutant shows additive effect (Fig 4B). We further calculated the fitness of each single mutant and double mutant to test whether K168A mutant is really additive to these two pathway or have genetic interaction with them followed the calculation method in previous study(35). As a result, K168A-rad5 double mutant have alleviated DNA damage fitness than estimated fitness which calculated based on single mutant fitness, indicates that K168A mutant have alleviating genetic interaction with RAD5 related DDT pathway, while K168A-rev3 double mutant had exactly the additive effect which indicates K168A has no genetic interaction between TLS pathway (Fig 4C). Moreover, considering that the damage sensitivity of K168A is highly resemble with *rad5△*and both Rad5p and PCNA play vital roles in template switching (TS) pathway, the critical DDT process is probably TS pathway. To test this hypothesis, we examined whether overexpression of TS-related genes, either upstream (*RAD51* related genes) or downstream (*SMC5*, *SMC6*, *RAD5*, *MMS2*), were able to suppress the damage sensitivity in K168A. Intriguingly, among all these genes, only overexpression of *RAD5* could rescue the damage sensitivity of K168A mutant to that of near wild type (Fig 4D), suggesting a RAD5-specific function might be involved. Since Rad5p is known to interact with PCNA (33), one possible explanation of this observation is that K168A weakens the interaction between Rad5p and PCNA. Therefore, we performed both the yeast two-hybrid analysis (Y2H) and co-immunoprecipitation (Co-IP) to monitor interactions between Rad5p and K168A or wild-type PCNA. As shown in Fig 4E-F, Rad5p is able to interact with the wild type PCNA and the interaction is greatly reduced in K168A mutant. Taking this together, the severe DNA damage sensitivity of K168A mutant might be a consequence of compromised interaction between Rad5p and PCNA.

### Interaction between PCNA and Rad5p is vital for Rad5p function implementation

Rad5p is a multi-functional protein involved in DNA damage tolerance (11, 36–40). To identify which function of Rad5p is specifically compromised in K168A, we constructed a series of Rad5p mutants including deletions and point mutations (Fig 5A). Over expression of most mutants cannot rescue damage sensitivity of K168A excepting the FN13,14AA (Fig 5B), a reported mutant deficient in the TLS pathway (41). On the other hand, over expression of ATPase and ubiquitin ligase defect mutants (DE681,682AA and CC914,917AA) somewhat enhanced the damage sensitivity of K168A (Fig 5B). Therefore, we showed that both the ubiquitin ligase function and ATPase function are essential for rescuing the damage sensitivity of K168A. Since both functions are closely related to the TS pathway (11), it is likely that the K168A mutant is deficient in the TS pathway through a Rad5-dependent mechanism. Since the point mutation which block the ubiquitin ligase function or ATPase function can also influence the helicase activity of RAD5, and the helicase mediated contribution to replication stress survival of RAD5 can be separable from TS pathway(42). To further confirm the replication stress defect of K168A is related to TS pathway, we tested whether K168A mutant could suppress the cold-sensitivity of *pol32△* cells, a phenotype which largely depends on PCNA ubiquitylation(43). The results show that K168A mutant could suppress the cold sensitivity of *pol32△*, to the same extent as rad5*△* mutation, which in consistent with our hypothesis that RAD5 related TS pathway is defected in K168A mutant.

**Fig 5.**
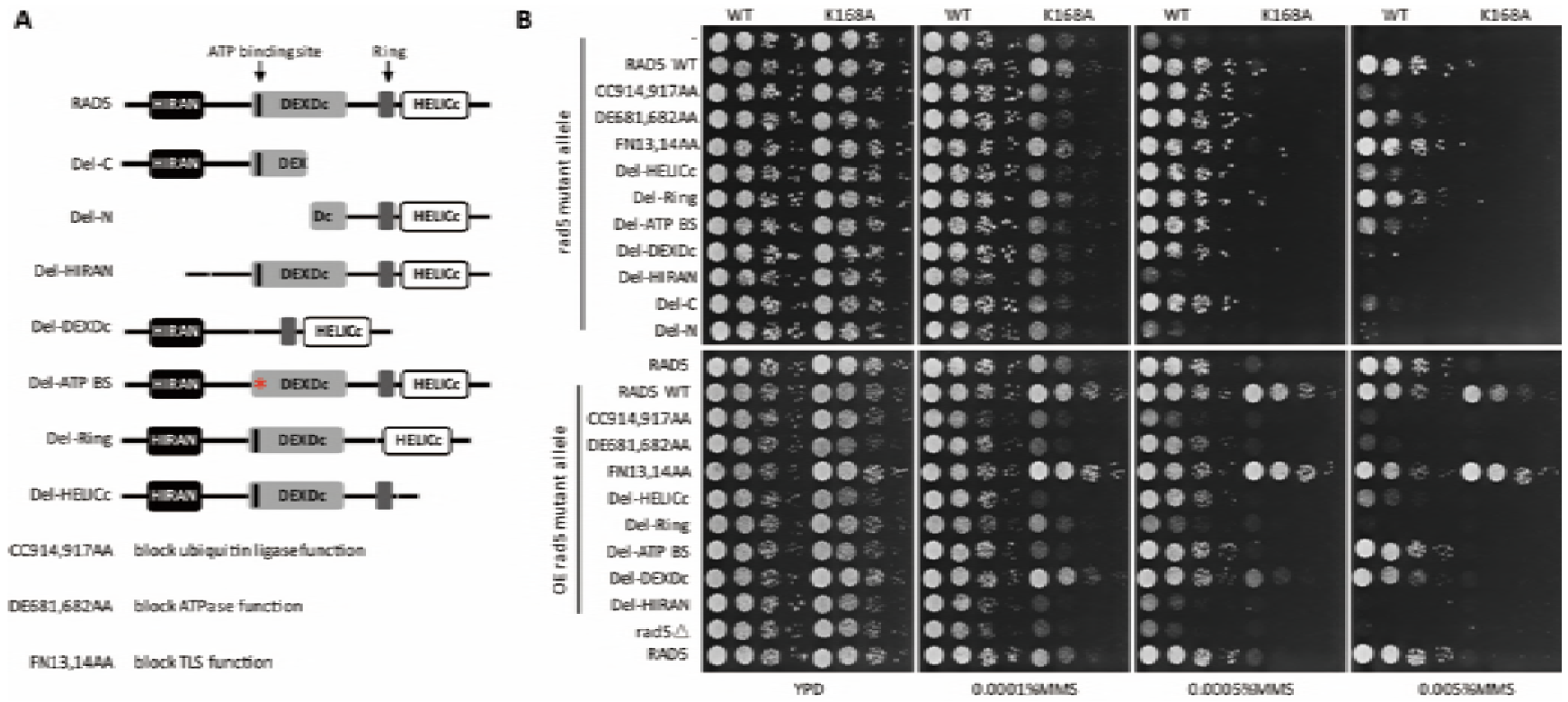
Both ubiquitin ligase and DNA helicase activity of RAD5 contribute to rescuing DDT deficicy of K168A. **(A)** Schematic diagram of protein domains and constructs of RAD5. **(B)** Tenfold serial dilutions of the indicated strains were spotted on medium supplemented with different amount of MMS. Overexpression of Rad5 is conducted by knocking in the GPD promoter to drive the expression of rad5 alleles.

**Fig 6.**
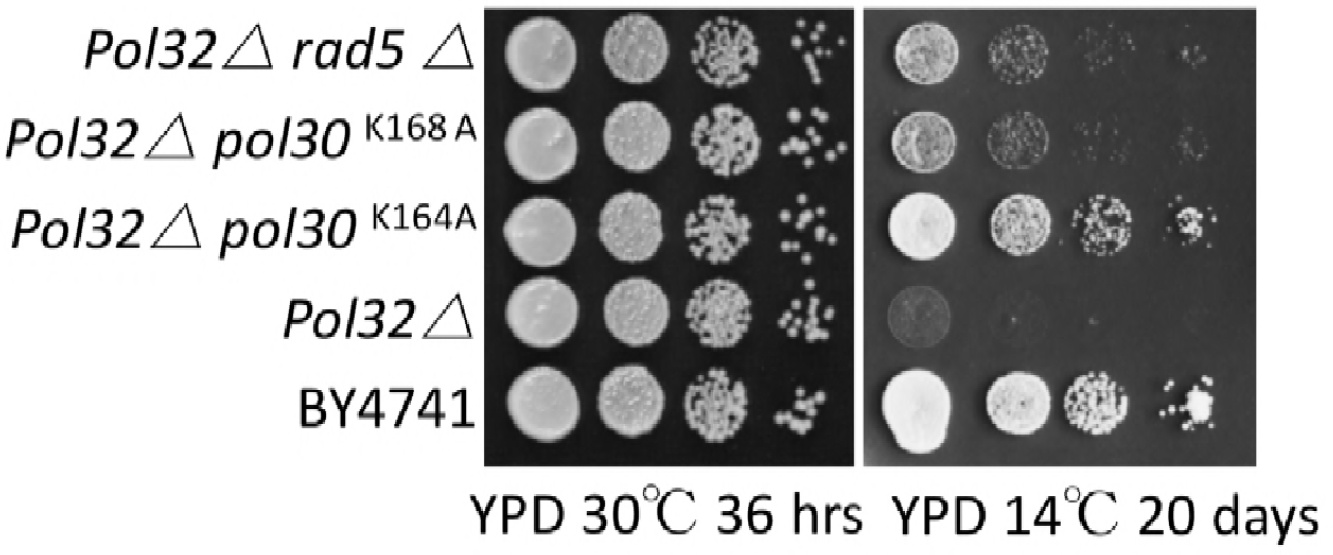
K168A mutant could suppress cold sensitivity of *pol32△*. Serial 1:10 dilutions of the indicated strains were spotted on two YPD plate to incubate at 30°C for 36 hrs or 14°C for 20 days.

Together, we revealed that the PCNA K168A mutant possesses severe sensitivity to DNA damage, and the deficiency does not result from compromising the post-translational modification at K164 but rather from reducing interaction with Rad5p.

## Discussion

In this study, a highly versatile library of PCNA mutants was carefully designed and chemically synthesized. Each mutant was integrated into the endogenous locus to generate a library of yeast each harboring one particular mutation, which allows us to systematically probe the contribution of each residue to many important functions of PCNA, such as the DNA damage response. New insights into the functional domains of PCNA were revealed.

### Crucial residues in PCNA for surviving genotoxic stress

Using the library, we identified a large number of mutants which are sensitive to DNA damage stresses caused by MMS, UV and HU. Interestingly, the DNA-damage-sensitive mutants are only sensitive to certain types of DNA-damaging reagents. Particularly, there were no mutants displayed sensitivity to CPT, a drug that can induce DNA damage by inhibiting topoisomerase I and hence causing DSBs (44). To confirm this result, we further tested the sensitivity of K168A and K164A, both of which are severely sensitive to HU, MMS and UV, to X-ray irradiation which also causes DSB. Neither of them showed elevated sensitivity to X-ray relative to the wild-type strain (data not shown). This result is consistent with the observation that these mutants are not sensitive to CPT, implying that PCNA may be dispensable for DSB repair.

After plotting the damage-sensitive mutants onto the crystal structure of PCNA trimer, we found these residues spread into several distinct regions, implying that different mechanisms might be applied. This is not unexpected given the fact that PCNA interacts with various partners to coordinate DNA replication and repair. The mutants in this library could potentially provide a resource to separate the multiple functions of PCNA by spatially disrupting interaction with a particular partner. Most functions of PCNA revealed previously are from the perspective of partner proteins, leading to more attention on C terminal and IDCL regions where are the hub sites for PIP-box targeting (4). However, in our screening, we identified more functional sites for DNA damage tolerance on the N terminal of PCNA, although the IDCL and nearby regions are also important. This N terminal make it possible for PCNA to partake some tasks from a previously reported “interaction hub”.

### The side chain of K168 correlates with DDT pathway by interacting with Rad5p

On account of our systematic mutagenesis principle, we can evaluate the importance of each residue of PCNA for DNA damage tolerance. A new mutant, K168A, attracted our attention for it had the most severe sensitivity to DNA damage among the alanine screening mutants and it showed much higher persistence and intensity of γH2A and RAD52 foci. We also demonstrated that the DNA damage sensitivity of this mutation is not caused by changing the SUMOylation or ubiquitination on PCNA. Actually, previous reports showed that both K164A and K168A mutants have no much deficiency in stimulating RFC ATPase or pol δ DNA synthesis (45). Although it is reported that PCNA ISGylation at K168 in a human cell line plays a crucial role in TLS termination (46), this kind of modification has not been reported in yeast yet. Our results inferred that K168A could block the function of Rad5p since K168A mutant drastically reduces the affinity to Rad5p while having no defect in interacting with other critical partners we have tested, such as ECO1, DPB2, RFC3, RAD18, and SRS2. (Fig S6). Indeed overexpression of Rad5p can rescue the DNA damage sensitivity of K168A mutant, consistent with a defect in Rad5 binding underlying the damage sensitivity phenotype. For futher consideration of the influence of K168A to TLS, we noticed that this mutant also disordered the affinities between PCNA and TLS polymerases. K168A decreased the affinity to RAD30 while increased the affinity to REV1 (Fig S6). This affinity disorder might explains why overexpression of RAD5 cannot rescue the K168A mutant to completely healthy status. In general, Rad5p is a multi-function protein with both ubiquitin E3 ligase activity and helicase activity (9, 11, 38, 40). Previous study showed clearly that this protein has important roles in error free leision bypass by promoting fork regression or template switching (40). Our domain dissection of Rad5p indicates that both the helicase and ubiquitin ligase functions of Rad5p are essential for rescuing the defects of the K168A mutant when Rad5 is overexpressed. Therefore, the interaction between Rad5p and PCNA is crucial for Rad5p to excecute its function.

Among the three known DDT pathways, TS pathway remains enigmatic since how PCNA poly-ubiquitination promotes this error free pathway is still under investigation (47, 48). Recently some efforts have been made to identify the factors correlated to PCNA poly-ubiquitination and TS pathway (49), which suggest that 9-1-1 complex might play a role in this pathway. However, the detailed mechanism remains to be elucidated (47, 49). Rad5p is known as a very important player in DDT pathway and the damage hyper sensitivity of *rad5* deletion strain emphasized the predominance of *RAD5* for lesion bypass (33). Since Rad5p is a large protein and there is still no complex structure of Rad5p binding to PCNA, we don’t know at a biophysical levelhow Rad5p interacts with PCNA. Here we showed that at least K168 is one of the key residues to mediate the interaction between PCNA and Rad5p, which might be vital for *RAD5*-dependent DDT pathway. Rad5p can also interact with Rad18p and Ubc13p which both dedicate to polyubiquitination of PCNA (50, 51). Our results indicate that the polyubiquitination of PCNA does not relay on direct interaction between PCNA and Rad5p, nevertheless the interaction is still vital for leision bypass. This may indicate that the interaction between PCNA and Rad5p can recruit Rad5p to proper site where Rad5p carry out its function, especially the helicase function which closely related to error free DDT pathway. Besides, the ubiquitin ligase function of Rad5p is also essential for rescuing damage sensitivity of K168A when it was overexpressed. This demonstrates that the synergistic effect of *RAD5* overexpression and K168A rely on PCNA polyubiquitination. However, whether the binding of Rad5p to PCNA is regulated by PCNA modification or whether this interaction has activity induction for Rad5p remain to be investigated. In all, this library of mutants could become a powerful tool to study the complicated genetic interactions between PCNA and other key factors to shed light on the regulation of DNA metablism.

## Material and methods

### Strains and antibodies

Assay specific strains were constructed with genotype listed in table S5. The reporter strain used for PCNA library construction and screening is QJY001. Antibodies used were anti-HA (Sigma-Aldrich, H3663), anti-PCNA (Abcam, 5E6/2), and H3 (Abcam, ab1791), anti-SMT3 (Abnova, MAB2093).

### Construction of the bacterial and yeast *POL30* library

The reporter strain (QJY001) is derived from S288C. The base construct was synthesized and cloned into *pRS414*. Fusion PCR was used to generate each individual mutant with unique barcode, for forward primer carry point mutation while corresponding reverse primer carry 20 nucleotide flapped sequence as barcode. All of the mutant plasmids were sequenced and supplied as bacterial stocks. To generate the yeast strains with integrated PCNA mutant, plasmids were digested with *Ear*I and transformed into QJY001 using standard lithium acetate protocol. The Leu+ Trp-G418^s^ transformants were subjected to colony PCR confirmation. The primers used were JDO236: GCAACCAAAGGAACCAAAGA; KanB: CTGCAGCGAGGAGCCGTAAT; KanC3: CCTCGACATCATCTGCCCAGA; JDO265: CAGGCATCGAATGGAAACTT; JDO264:AAGCCTCCTCGAACTTAGCC; JQO44: ATTGACAAGGAGGAGGGCACC; Two independent colonies for each mutant were obtained and subjected to plasmid shuffling assay as well as to generate the yeast PCNA mutant library.

### Phenotypic growth assays

The yeast mutant stains were arrayed on the rectangle SC–Leu plates. Similar amount of cells for each mutant were suspended in 100 μL ddH_2_O, spotted on SC–Leu plate and grown at 30°C overnight. Then the cells were diluted to density at A_600_=0.1 for 3 times to generate 4 concentration gradients (10-fold dilution). Finally, 3 μL of cell suspension were spotted on different media and incubated accordingly.

For DNA damage sensitivity test, YPD plates were supplemented with 200 mM hydroxyurea (HU), 0.005% 0.015% or 0.03% methylmethanesulonate (MMS) or SC–Leu plate supplemented with 8 μ g/mL camptothecin. To test UV sensitivity, cells were dropped on YPD plates, exposed to different dosage of UV irradiation (50 J/m^2^ 80 J/m^2^ and 100 J/m^2^), wrapped with foil and incubated at 30°C. Benomyl sensitivity was tested on YPD medium containing 10 μg/mL benomyl. For temperature sensitivity, cells were dropped onto YPD plates and incubated at 16°C, 37°C or 39°C.

### Structure mapping

The mutated residues were highlighted in PCNA crystal structure using Pymol. The PDB number of each structure was shown in the figure legend.

### Prepare the whole cell protein lysate

Cells were grown overnight in YPD or appropriate synthetic media at 30°C, diluted to A_600_=0.1 in 5 mL fresh media and grown to final A_600_=0.8-1.0. After the required treatments, cells were harvested and resuspended in water to obtain the same amount of cells for each sample. Whole cells lysates were prepared by resuspending the cell pellets in 100 μL ddH_2_O, adding 100 μL 0.2 M NaOH, vortexing the mix briefly and staying at RT for 5 min. Cells were pelleted and resuspend in 1×SDS loading buffer (0.0625 M Tris-HCl pH6.8, 2% SDS, 10% glycerol, 0.003% bromophenol blue, 5% β-Mercaptoethanol). The resuspended samples were boiled at 100°C for 5 min, transferred to ice immediately, kept for 2 min and then centrifugated at 14000 g for 2 min. The supernatant was used as the whole cell lysate.

### Cell cycle analysis

The logarithmic cells were arrested at G1 by 10 μM of α-factor for 3 hrs, cells were washed and resuspended in fresh medium with 50 μg/mL pronase. Cells that under normal growth condition were collected every 15 min whereas cells that under MMS treatment were collected every 30 min. Cells were fixed, stained and anlyazed by 70% ethanol, PI (Propidium Iodide) and BD FACSCalibur respectively.

### Yeast two-hybrid assay

Wild-type or K168A mutant *PCNA* was cloned into standard DNA binding domain (BD) plasmid and the candidate genes were cloned into activating domain (AD) plasmid. AD and BD plasmids were co-transformed into reporter strain JDY26 and at least two independent transformants were tested. Cells were dropped on SC–Leu–Trp and SC–Leu–Trp–His medium.

### Cell growth curve

BioTek™ Epoch™ 2 Microplate Spectrophotometer was used for growth curve determination. Each strain (A_600_=0.1) was cultivated in 384-well microplates containing 40 μL indicated medium. Cell growth was monitored by measuring A_600_ every 15 min. Optical densities (OD) were corrected by a calibration function to convert the recoded OD to real OD. The calibration function is as following:

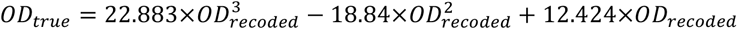

### Classification of genetic interaction

The genetic interaction were tested and calculated as previously described(35). In brief, BioTek™ Epoch™ 2 Microplate Spectrophotometer was used for determine the doubling time (*D*) of each strain and the fitness *(W)* of a mutation strain *x* was defined as the ratio of the *D* value of the wild-type to the mutation strain (*W*=*D*_*wt*_/*D*_*x*_). Fitness of double mutant was predicted for non-interacting gene pair (*W*_*xy*_=*W*_*x*_ × *W*_*y*_) under the multiplicative model. The genetic interaction was classified with Z-test of estimated fitness and the real fitness of double mutant.

### Yeast protein purification

His-tagged proteins were enriched by Nickle-NTA beads in a denatured way as reported previously (52). Equal amount of logarithmic cells were harvested and treated with 0.1 M NaOH for 10 min at room temperature. Cells were resuspended in 250 μL of lysis buffer (60 mM Tris-HCl pH 6.8, 5% glycerol, 2% SDS, 4% 2-mercaptoethanol) and boiled at 100°C for 8 min. Samples were centrifuged with 1500 g for 25 min and 200 μL of supernatant was transferred into a new tube containing 750 μL of dilution buffer (10 mM Tris-HCl pH8, 100 mM NaPhosphate pH 8, 8 M Urea) and 50 μL of Nickle-NTA beads. Samples were incubated at 4°C for 2 hrs on rotator and then washed with dilution buffer for three times and then with wash buffer (10 mM NaPhosphate pH 8.0, 500 mM NaCl, 0.5% NP-40, 10 mM Imidazole) for three times. His-tagged protein was eluted with elution buffer (10 mM NaPhosphate pH 8.0, 500 mM NaCl, 0.5% NP-40, 400 mM imidazole) by incubating at 4°C for 1 hr and then centrifuged to pellet the beads. Supernatant was taken out for western blot analysis.

### Yeast GST-protein Co-IP

Overnight yeast culture was diluted to A_600_=0.2 with 20 mL fresh medium and cultured at 30° C with vigorous shaking until the cell density reach to A_600_=1.2. Same amount of cells were harvested and washed with ice-cold water, and then resuspended in 500 μL of breaking buffer (TNT buffer with 1 mM DTT, protease inhibitor cocktail, 1 mM PMSF). 250 μL of glass beads were added into the tube, which was shaken at 1500 g for 1min. Samples were then transferred on ice for 2 min. Repeated the shaking procedure for 6 times. Samples were centrifuged with 13500 g for 5 min at 4°C and the supernatant were transferred to new tubes as whole cell extraction. Same amount of whole cell extractions were added to 50 μL of PBS and GSH beads, which was prewashed by TNT buffer (50 mM Tris-HCl pH 7.5, 50 mM NaCl, 0.1% TritonX-100, 0.5 mM EDTA, 10% glycerol). Samples were incubated at 4 °C for 3 hours and then washed by 500 μL of TNT butter for three times. CO-IP sample was eluted with 50 μL of 15 mM GSH-TNT buffer and the supernatant was taken for western blot analysis.

### Denaturing IP

IP for detection PCNA modification was exactly followed lab protocol of Tansey (*http://www.tanseylab.com/the-myc-family-of-oncoprote/denat_ip_yeast.pdf*).

## Acknowledgment

This work was supported by the National Key Research and Development Program of China (Grant 2017YFA0505103), by the National Natural Science Foundation of China (31725002), by Bureau of International Cooperation, Chinese Academy of Sciences (172644KYSB20170042) and by the Key Research Program of the Chinese Academy of Science (KFZD-SW-215).

## Reference

1. Hoeijmakers JHJ. 2009. DNA damage, aging, and cancer. N Engl J Med 361:1475–1485.

2. Kurth I, O’Donnell M. 2013. New insights into replisome fluidity during chromosome replication. Trends in Biochemical Sciences 38:195–203.

3. Stoimenov I, Helleday T. 2009. PCNA on the crossroad of cancer. Biochemical Society Transactions 37:605–613.

4. Moldovan G-L, Pfander B, Jentsch S. 2007. PCNA, the Maestro of the Replication Fork. Cell 129:665–679.

5. Chang DJ, Cimprich KA. 2009. DNA damage tolerance: when it’s OK to make mistakes. Nature Chemical Biology 5:82–90.

6. Branzei D, Szakal B. 2016. DNA damage tolerance by recombination: Molecular pathways and DNA structures. DNA Repair 44:68–75.

7. Freudenthal BD, Gakhar L, Ramaswamy S, Washington MT. 2010. Structure of monoubiquitinated PCNA and implications for translesion synthesis and DNA polymerase exchange. Nat Struct Mol Biol 17:479–484.

8. Boiteux S, Jinks-Robertson S. 2013. DNA repair mechanisms and the bypass of DNA damage in Saccharomyces cerevisiae. Genetics 193:1025–1064.

9. Giannattasio M, Zwicky K, Follonier C, Foiani M, Lopes M, Branzei D. 2014. Visualization of recombination-mediated damage bypass by template switching. Nature Structural & Molecular Biology 21:884–892.

10. Vanoli F, Fumasoni M, Szakal B, Maloisel L, Branzei D. 2010. Replication and recombination factors contributing to recombination-dependent bypass of DNA lesions by template switch. PLoS Genet 6:e1001205.

11. Minca EC, Kowalski D. 2010. Multiple Rad5 activities mediate sister chromatid recombination to bypass DNA damage at stalled replication forks. Mol Cell 38:649–661.

12. Pfander B, Moldovan G-L, Sacher M, Hoege C, Jentsch S. 2005. SUMO-modified PCNA recruits Srs2 to prevent recombination during S phase. Nature 436:428–433.

13. Streich Jr FC, Lima CD. 2016. Capturing a substrate in an activated RING E3/E2-SUMO complex. Nature 536:304–308.

14. Lau WCY, Li Y, Zhang Q, Huen MSY. 2015. Molecular architecture of the Ub-PCNA/Pol η complex bound to DNA. Sci Rep 5.

15. Ayyagari R, Impellizzeri KJ, Yoder BL, Gary SL, Burgers PM. 1995. A mutational analysis of the yeast proliferating cell nuclear antigen indicates distinct roles in DNA replication and DNA repair. Mol Cell Biol 15:4420–4429.

16. Amin NS, Holm C. 1996. In vivo analysis reveals that the interdomain region of the yeast proliferating cell nuclear antigen is important for DNA replication and DNA repair. Genetics 144:479–493.

17. Eissenberg JC, Ayyagari R, Gomes XV, Burgers PM. 1997. Mutations in yeast proliferating cell nuclear antigen define distinct sites for interaction with DNA polymerase delta and DNA polymerase epsilon. Mol Cell Biol 17:6367–6378.

18. Chen C, Merrill BJ, Lau PJ, Holm C, Kolodner RD. 1999. Saccharomyces cerevisiae pol30 (proliferating cell nuclear antigen) mutations impair replication fidelity and mismatch repair. Molecular and cellular biology 19:7801–7815.

19. Zamir L, Zaretsky M, Fridman Y, Ner-Gaon H, Rubin E, Aharoni A. 2012. Tight coevolution of proliferating cell nuclear antigen (PCNA)-partner interaction networks in fungi leads to interspecies network incompatibility. Proceedings of the National Academy of Sciences 109:E406–E414.

20. Goellner EM, Smith CE, Campbell CS, Hombauer H, Desai A, Putnam CD, Kolodner RD. 2014. PCNA and Msh2-Msh6 Activate an Mlh1-Pms1 Endonuclease Pathway Required for Exo1-Independent Mismatch Repair. Molecular Cell 55:291–304.

21. Zhang Z, Shibahara K, Stillman B. 2000. PCNA connects DNA replication to epigenetic inheritance in yeast. Nature 408:221–225.

22. Miller A, Chen J, Takasuka TE, Jacobi JL, Kaufman PD, Irudayaraj JMK, Kirchmaier AL. 2010. Proliferating Cell Nuclear Antigen (PCNA) Is Required for Cell Cycle-regulated Silent Chromatin on Replicated and Nonreplicated Genes. Journal of Biological Chemistry 285:35142–35154.

23. Dieckman LM, Washington MT. 2013. PCNA trimer instability inhibits translesion synthesis by DNA polymerase η and by DNA polymerase δ. DNA repair 12:367–376.

24. Hishiki A, Shimizu T, Serizawa A, Ohmori H, Sato M, Hashimoto H. 2008. Crystallographic study of G178S mutant of human proliferating cell nuclear antigen. Acta Crystallographica Section F Structural Biology and Crystallization Communications 64:819–821.

25. Kondratick CM, Boehm EM, Dieckman LM, Powers KT, Sanchez JC, Mueting SR, Washington MT. 2016. Identification of New Mutations at the PCNA Subunit Interface that Block Translesion Synthesis. PLOS ONE 11:e0157023.

26. Dieckman LM, Freudenthal BD, Washington MT. 2012. PCNA Structure and Function: Insights from Structures of PCNA Complexes and Post-translationally Modified PCNA, p. 281–299. In MacNeill, S (ed.), The Eukaryotic Replisome: a Guide to Protein Structure and Function. Springer Netherlands, Dordrecht.

27. Winzeler EA. 1999. Functional Characterization of the S. cerevisiae Genome by Gene Deletion and Parallel Analysis. Science 285:901–906.

28. Ortega J, Li JY, Lee S, Tong D, Gu L, Li G-M. 2015. Phosphorylation of PCNA by EGFR inhibits mismatch repair and promotes misincorporation during DNA synthesis. Proc Natl Acad Sci USA 112:5667–5672.

29. Wang S-C, Nakajima Y, Yu Y-L, Xia W, Chen C-T, Yang C-C, McIntush EW, Li L-Y, Hawke DH, Kobayashi R, Hung M-C. 2006. Tyrosine phosphorylation controls PCNA function through protein stability. Nature Cell Biology 8:1359–1368.

30. Dai J, Hyland EM, Yuan DS, Huang H, Bader JS, Boeke JD. 2008. Probing Nucleosome Function: A Highly Versatile Library of Synthetic Histone H3 and H4 Mutants. Cell 134:1066–1078.

31. McNally R, Bowman GD, Goedken ER, O’Donnell M, Kuriyan J. 2010. Analysis of the role of PCNA-DNA contacts during clamp loading. BMC Structural Biology 10:3.

32. Vijayakumar S, Chapados BR, Schmidt KH, Kolodner RD, Tainer JA, Tomkinson AE. 2007. The C-terminal domain of yeast PCNA is required for physical and functional interactions with Cdc9 DNA ligase. Nucleic Acids Res 35:1624–1637.

33. Hoege C, Pfander B, Moldovan G-L, Pyrowolakis G, Jentsch S. 2002. RAD6-dependent DNA repair is linked to modification of PCNA by ubiquitin and SUMO. Nature 419:135–141.

34. Papouli E, Chen S, Davies AA, Huttner D, Krejci L, Sung P, Ulrich HD. 2005. Crosstalk between SUMO and ubiquitin on PCNA is mediated by recruitment of the helicase Srs2p. Molecular cell 19:123–133.

35. Onge RPS, Mani R, Oh J, Proctor M, Fung E, Davis RW, Nislow C, Roth FP, Giaever G. 2007. Systematic pathway analysis using high-resolution fitness profiling of combinatorial gene deletions. Nature Genetics 39:199–206.

36. Ahne F, Jha B, Eckardt-Schupp F. 1997. The RAD5 gene product is involved in the avoidance of non-homologous end-joining of DNA double strand breaks in the yeast Saccharomyces cerevisiae. Nucleic Acids Res 25:743–749.

37. Chen S, Davies AA, Sagan D, Ulrich HD. 2005. The RING finger ATPase Rad5p of Saccharomyces cerevisiae contributes to DNA double-strand break repair in a ubiquitin-independent manner. Nucleic Acids Res 33:5878–5886.

38. Gangavarapu V, Haracska L, Unk I, Johnson RE, Prakash S, Prakash L. 2006. Mms2-Ubc13-dependent and-independent roles of Rad5 ubiquitin ligase in postreplication repair and translesion DNA synthesis in Saccharomyces cerevisiae. Mol Cell Biol 26:7783–7790.

39. Pagès V, Bresson A, Acharya N, Prakash S, Fuchs RP, Prakash L. 2008. Requirement of Rad5 for DNA polymerase zeta-dependent translesion synthesis in Saccharomyces cerevisiae. Genetics 180:73–82.

40. Blastyák A, Pintér L, Unk I, Prakash L, Prakash S, Haracska L. 2007. Yeast Rad5 Protein Required for Postreplication Repair Has a DNA Helicase Activity Specific for Replication Fork Regression. Molecular Cell 28:167–175.

41. Xu X, Lin A, Zhou C, Blackwell SR, Zhang Y, Wang Z, Feng Q, Guan R, Hanna MD, Chen Z, Xiao W. 2016. Involvement of budding yeast Rad5 in translesion DNA synthesis through physical interaction with Rev1. Nucl Acids Res gkw183.

42. Choi K, Batke S, Szakal B, Lowther J, Hao F, Sarangi P, Branzei D, Ulrich HD, Zhao X. 2015. Concerted and differential actions of two enzymatic domains underlie Rad5 contributions to DNA damage tolerance. Nucleic Acids Res 43:2666–2677.

43. Karras GI, Jentsch S. 2010. The RAD6 DNA Damage Tolerance Pathway Operates Uncoupled from the Replication Fork and Is Functional Beyond S Phase. Cell 141:255–267.

44. Pommier Y, Redon C, Rao VA, Seiler JA, Sordet O, Takemura H, Antony S, Meng L, Liao Z, Kohlhagen G, Zhang H, Kohn KW. 2003. Repair of and checkpoint response to topoisomerase I-mediated DNA damage. Mutation Research/Fundamental and Molecular Mechanisms of Mutagenesis 532:173–203.

45. Fukuda K, Morioka H, Imajou S, Ikeda S, Ohtsuka E, Tsurimoto T. 1995. Structure-function relationship of the eukaryotic DNA replication factor, proliferating cell nuclear antigen. J Biol Chem 270:22527–22534.

46. Park JM, Yang SW, Yu KR, Ka SH, Lee SW, Seol JH, Jeon YJ, Chung CH. 2014. Modification of PCNA by ISG15 Plays a Crucial Role in Termination of Error-Prone Translesion DNA Synthesis. Molecular Cell 54:626–638.

47. Choe KN, Moldovan G-L. 2017. Forging Ahead through Darkness: PCNA, Still the Principal Conductor at the Replication Fork. Molecular Cell 65:380–392.

48. 2012. The Eukaryotic Replisome: a Guide to Protein Structure and Function. Springer Netherlands, Dordrecht.

49. Karras GI, Fumasoni M, Sienski G, Vanoli F, Branzei D, Jentsch S. 2013. Noncanonical role of the 9-1-1 clamp in the error-free DNA damage tolerance pathway. Mol Cell 49:536–546.

50. Ulrich HD, Jentsch S. 2000. Two RING finger proteins mediate cooperation between ubiquitin - conjugating enzymes in DNA repair. The EMBO Journal 19.

51. Ball LG, Xu X, Blackwell S, Hanna MD, Lambrecht AD, Xiao W. 2014. The Rad5 helicase activity is dispensable for error-free DNA post-replication repair. DNA Repair (Amst) 16:74–83.

52. Masumoto H, Hawke D, Kobayashi R, Verreault A. 2005. A role for cell-cycle-regulated histone H3 lysine 56 acetylation in the DNA damage response. Nature 436:294–298.

